# The world’s first cloned golden wild yak via interspecific SCNT: 4800m donor origin and 4200m vitrified blastocyst transfer

**DOI:** 10.64898/2026.03.29.715085

**Authors:** Dawei Yu, Qiang Zhang, Lei Cao, Shigang Gu, Yurong Zhang, Chang Liu, Kangjun Yin, Jing Wang, Bo Pan, Yang Liu, Guangbin Zhou, Daoliang Lan, Yongye Huang, Wangdui Basang

## Abstract

Primarily inhabiting the harsh, high-altitude environment of the Qiangtang National Nature Reserve exceeding 5,000 meters above the sea (m.a.s.l.), the golden wild yak is critically endangered, with fewer than 300 individuals remaining in the world, a situation exacerbated by the significant challenges of conducting research and conservation of their genetic resources. Somatic cell nuclear transfer (SCNT) can be an effective method for their preservation, but facing several obstacles in this context, including the hypoxic stress at high altitude that impairs embryonic development due to *in vitro* manipulation, and constraints of long-distance embryo transport. In the present study, the ear tissue was collected from a childhood male golden wild yak at Xizang Geye Wildlife Rescue Station (4800 m.a.s.l.) and send to Institute of Animal Science at Beijing to derive fibroblast cells. Using fibroblast cells of the golden wild yak as nuclear donors, and bovine oocytes from a local slaughterhouse at Beijing as recipients, the interspecific SCNT (iSCNT) embryos were generated and *in vitro* developed to blastocysts. To maintain the embryonic viability after long-distance transportation from Beijing to Xizang, iSCNT blastocysts were subjected to cryopreservation by vitrification method. Thawing of vitrified iSCNT blastocysts were completed at Xizang Dangxiong Yak Breeding Innovation Base (4200 m.a.s.l.), and transferred into the uterine horn of domestic yaks. 257 days after blastocyst transfer, a cloned golden wild yak was successfully harvested on January 10, 2026. This work demonstrates, for the first time, that interspecies somatic cell nuclear transfer can successfully produce a cloned offspring under extreme conditions, spanning 4800 m.a.s.l. donor origin, long-distance vitrified embryo transportation, and high-altitude blastocyst transfer at 4200 m.a.s.l., establishing a viable strategy for conserving critically endangered high-altitude species.

## 1. Introduction

Wild yak, as a wildlife species in the Qinghai-Xizang plateau, possesses significant ecological value and economic potential ^[1-3]^. However, due to climate change, habitat fragmentation, and human activities, the remaining wild yak populations are classified as vulnerable by the International Union for Conservation of Nature and Natural Resources (IUCN) Red List ^[4, 5]^. The accelerating degradation of yak populations in alpine regions underscores the urgent need for germplasm conservation. However, difficulties in collecting live samples and artificially breeding by conventional methods further enhance their vulnerability.

The golden wild yak, with a global population of fewer than 300 individuals mainly inhabiting in Ritu county (4500 m.a.s.l.), Geji county (4800 m.a.s.l.) and Qiangtang National Nature Reserve (5000 m.a.s.l.), faces an imminent risk of extinction. According to the previous reports, there are fewer than 300 golden wild yaks. In recent years, assisted reproductive technologies (ARTs), particularly somatic cell nuclear transfer (SCNT), have provided new technical methods for the salvage conservation of endangered species ^[6]^. Although SCNT technology has been successfully applied to the propagation of various livestock and wild animals, its application is limited by the difficulty of obtaining oocytes from wild species. The emergence of iSCNT technology has provided new ideas for cloning endangered species. Currently, iSCNT technology has cloned multiple animals, including Felis silvestris lybica, Canis latrans, boars, and Pien-niu ^[7-10]^, verifying the significant potential of this technology in biological breeding and the conservation of genetic resources. Therefore, this study aims to achieve a breakthrough in golden wild yak through iSCNT, thereby providing a possibility for the long-term conservation of wild genetic resources in extreme environments.

## 2. Results

### 2.1 Establishment and assessment of donor cells from the golden wild yak ear tissue

Fibroblast cell lines were isolated and established from the living ear tissue. When these cells were subcultured to the third passage (P3), they exhibited typical fibroblast morphology (Figure 1A). The viability and proliferation capacity of donor cells were assessed using CCK-8 and flow cytometry. The results of the CCK-8 growth curve assay showed that this cell line exhibited stable cell proliferation trend (Figure 1B). Flow cytometry analysis further revealed that the total apoptosis rate of donor cells is approximately 11.04 % (Figure 1C). To confirm the genetic stability of this cell line, we performed chromosome karyotyping and G-band analysis on P5 cells (Figure 1D). The analysis results indicated that the chromosome number remained stable at 60, consisting of 29 pairs of autosomes and one pair of sex chromosomes (Figure 1E). The centromeres of all chromosomes were clearly distinguishable, without structural abnormalities observed. These results suggest that this cell line maintained excellent genetic stability during passaging.

**Figure 1.**
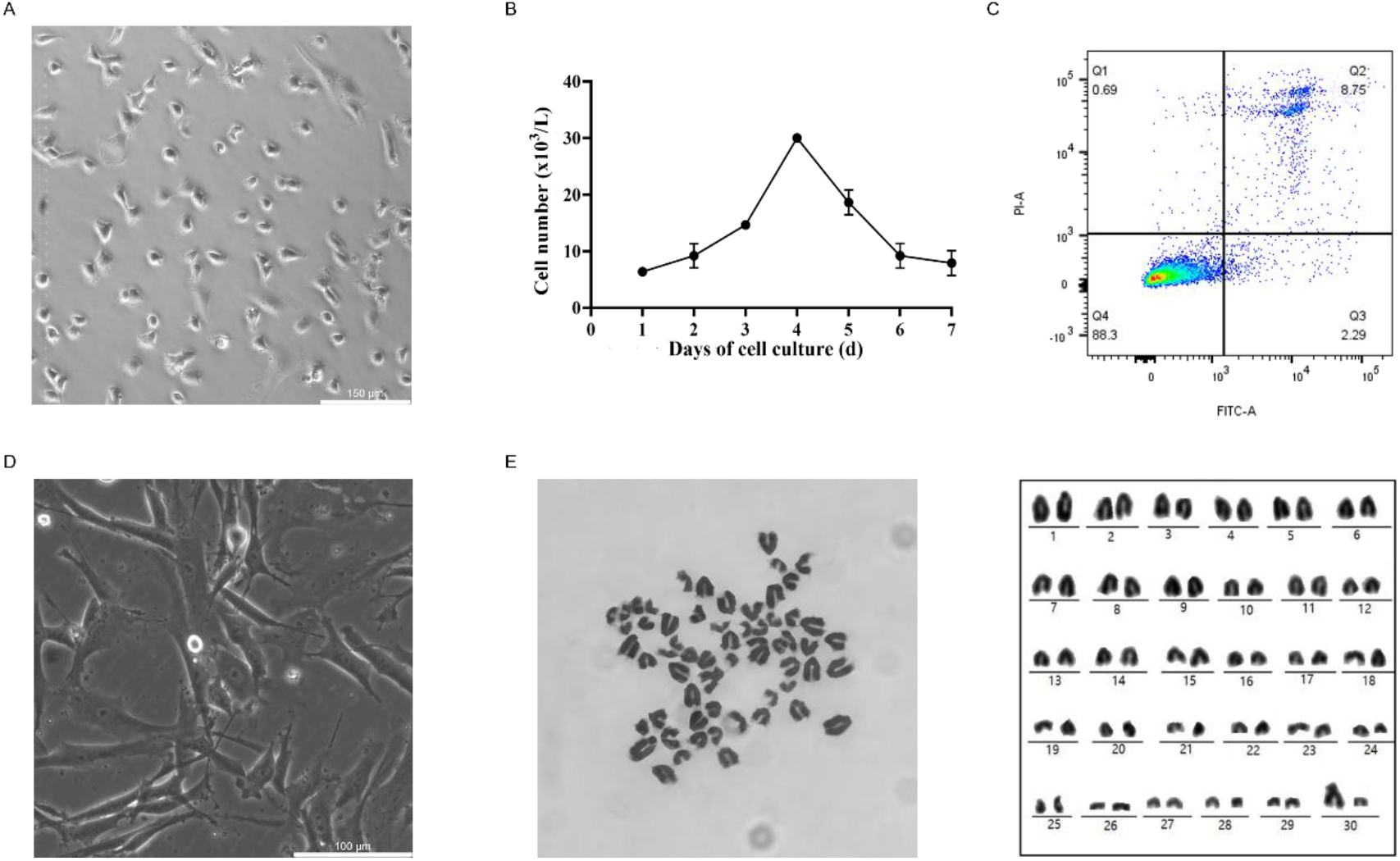
Establishment and assessment of donor cells from the golden wild yak ear tissue. A. Cell morphology of the third passage (P3) cells. Scale bars, 150 µm. B. The CCK-8 growth curve assay of donor cells. C. Flow cytometry measurement of apoptosis in donor cells. D. Growth morphology of the fifth passage (P5) cells. Scale bars, 100 µm. E. Karyotype (left side) and chromosome sequencing (right side) of P5 cells.

### 2.2 Construction of iSCNT embryos and evaluation of their developmental ability

The high-quality golden wild yak fibroblast cells were selected as nuclear donors and bovine oocytes from the local slaughterhouse as recipients to construct iSCNT embryos. Using bovine SCNT embryos and *in vitro* fertilization (IVF) embryos as control groups, the developmental ability of iSCNT embryos was assessed (Figure 2A). As shown in Table 1, the cleavage and blastocyst formation rates reached 81.62 ± 2.68 % and 15.39 ± 7.13 %, respectively. To further determine the developmental ability of iSCNT embryos, immunofluorescence staining was performed for trophectoderm cells (CDX2) and inner cell mass (SOX2) (Figure 2B). Results found that iSCNT embryos could differentiate normally to form the inner cell mass (ICM) and trophoblast (TE), suggesting that iSCNT embryos possess the totipotency to develop into complete individuals.

**Table 1.** Assessment of the developmental potential of iSCNT embryos.

| Donor cell | Repetition | Enucleated<br>oocyte | % cleaved ±<br>S.E.M. | % blastocyst ±<br>S.E.M. |
| --- | --- | --- | --- | --- |
| IVF | 3 | — | 94.00 ± 1.91 | 39.89 ± 5.49 |
| SCNT | 3 | 298 | 94.42 ± 2.30 | 37.43 ± 1.34 |
| iSCNT | 5 | 1125 | 81.62 ± 2.68 | 15.39 ± 7.13 |

**Figure 2.**
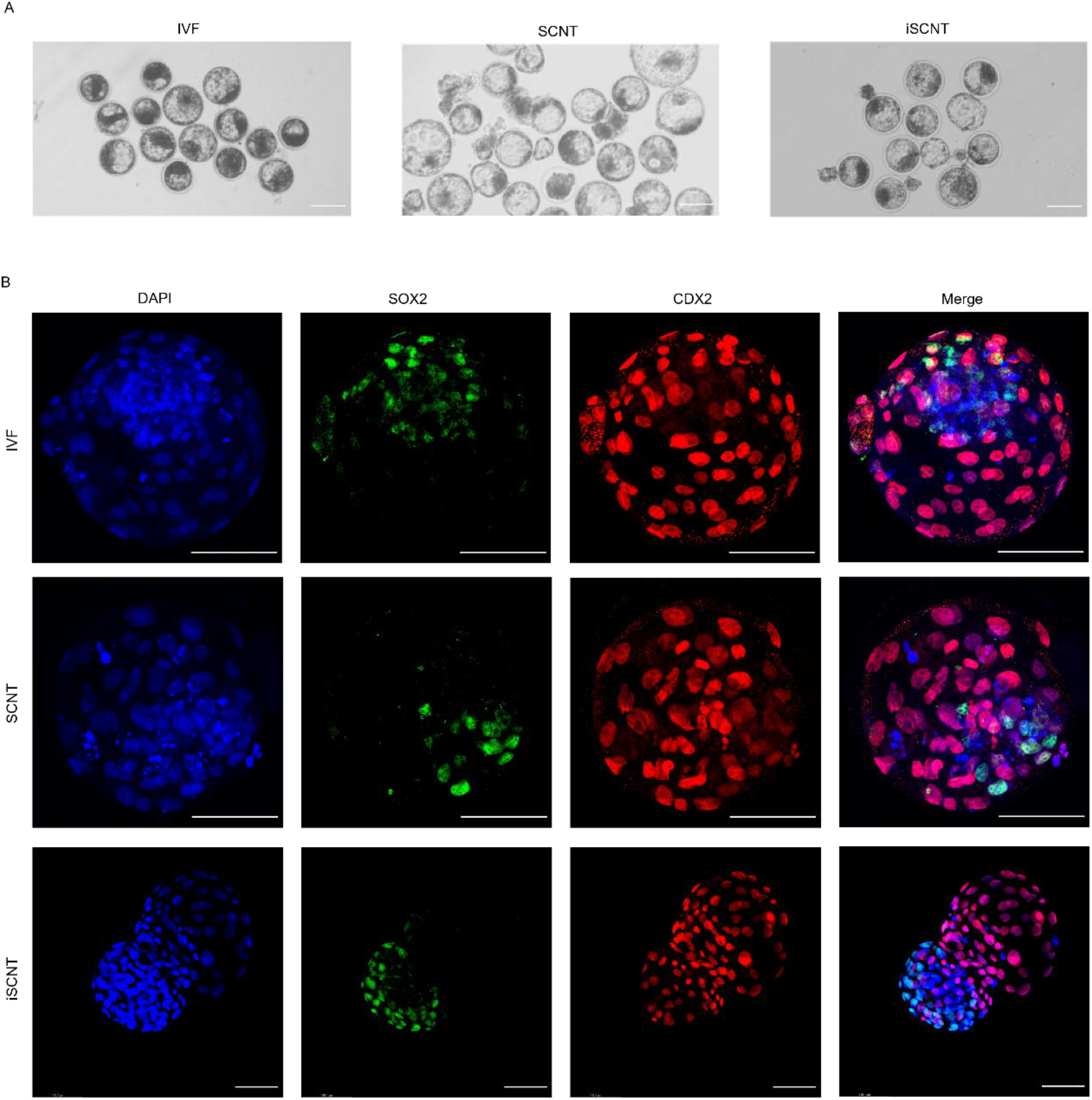
Construction of iSCNT embryos and evaluation of their developmental ability. A. Morphology of bovine IVF (left side), SCNT (middle side), and iSCNT blastocyst. (right side). Scale bar, 100μm. B. Immunofluorescence staining of trophoblast cells (CDX2) and inner cell mass (SOX2) in blastocyst. Scale bar, 100μm.

### 2.3 Birth of iSCNT golden wild yak

To complete the iSCNT embryo transfer, thirty-four domestic yaks underwent synchronized estrus treatment at the Xizang Dangxiong Yak Breeding Innovation Base (4200 m.a.s.l.). Twenty-one healthy domestic yaks with synchronized oestrus were selected as surrogate yaks, and well-developed recombinant embryos were transferred into the uteri of these surrogate yaks. Pregnancies were confirmed in two surrogate yaks via ultrasound examination, resulting in a 9.52 % pregnancy rate (Figure 3A). One of the surrogate yaks miscarried midway through gestation. On January 10, 2026, a live cloned golden wild yak was born, weighing 34.8 kilograms (Figure 3B). Apparently, the coat color of the newborn cloned golden wild yak was different from the surrogate yak (Figure 3C), but similar to the iSCNT donor cells provider (Figure 3D). To verify its genetic origin, short tandem repeat (STR) polymorphism analysis on the cloned golden wild yak, iSCNT donor cells provider and surrogate yak were conducted (Figure 3E). The STR results show that the microsatellite loci of the cloned golden yak were identical to those of the donor cells. Collectively, these results suggested that iSCNT technology can achieve the cloning and breeding of ultra-high-altitude endangered mammals. It provided a reliable pathway for the long-term preservation of golden wild yak and the restoration of its population.

**Figure 3.**
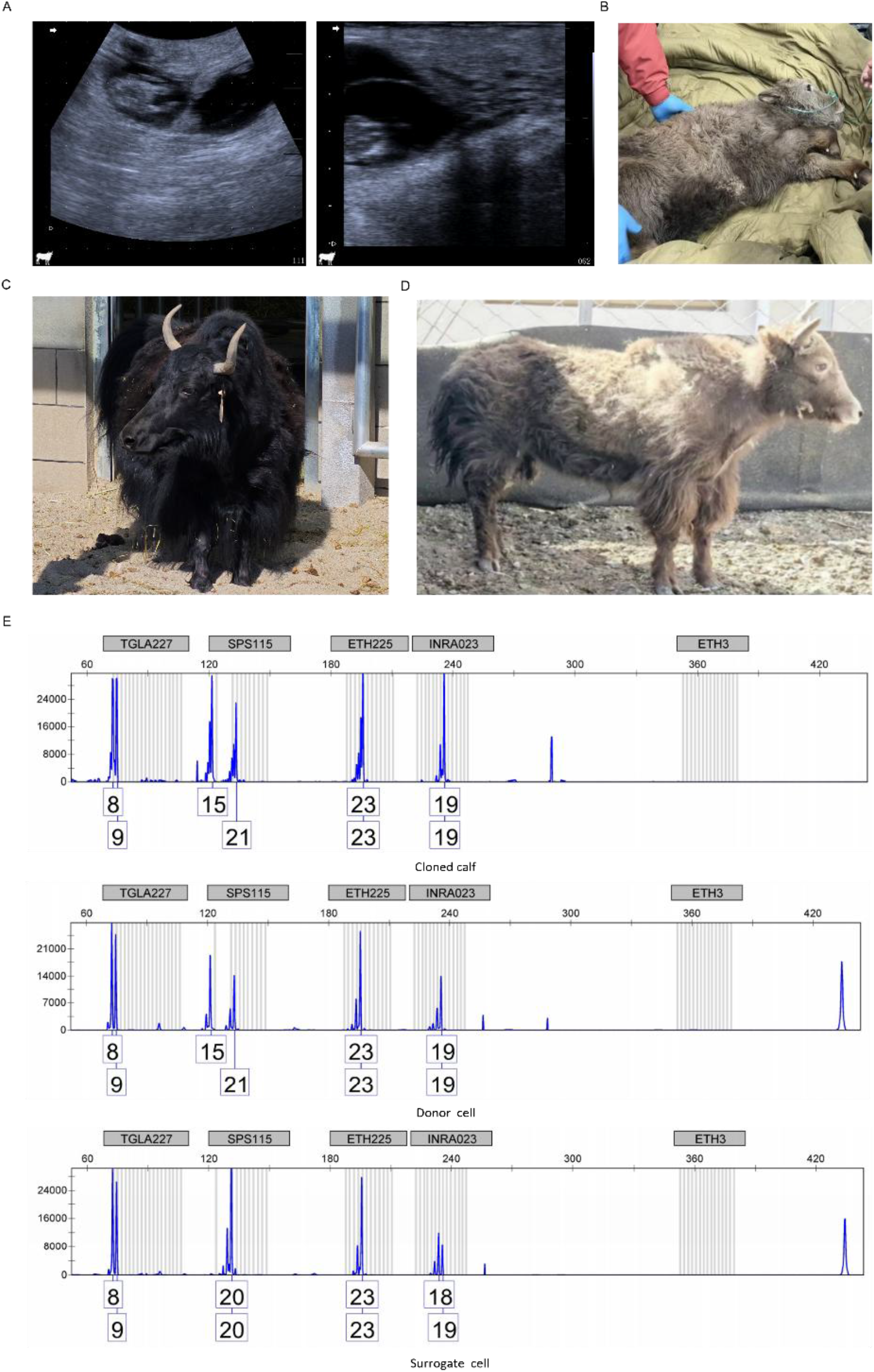
Birth of the iSCNT golden wild yak. (A) Pregnancies were confirmed in two surrogate yaks via ultrasound detection. (B) A live cloned golden wild yak was born, weighing 34.8 kilograms on January 10, 2026. (C) Surrogate yak for the cloned golden wild yak (B). (C) iSCNT donor cell provider. (E) Representative STR loci of the cloned golden wild yak (B), iSCNT donor cells provider (D), and surrogate yak (C).

## 3. Discussion

The present study had successfully produced the first live offspring of the golden wild yak via iSCNT, marking a crucial step forward in the conservation of endangered animals. As a key species in the Qinghai-Xizang Plateau ecosystem, the wild yak population has been declining due to environmental degradation and human activities, and listed as a vulnerable species ^[4]^. Although SCNT technology is becoming increasingly mature, its development and application are still limited by its low efficiency ^[11]^. SCNT has high requirements for technical skills, oocyte quality, and donor cell quality ^[12]^. Long-term *in vitro* culture might induce chromosomal aneuploidy in cells, and karyotypic abnormalities in donor cells are one of the main causes of developmental arrest or miscarriage in cloned embryos ^[13]^. Thus, we assessed the quality of donor cells through karyotype analysis. Our results suggest the donor cells were of good quality, with stable chromosome numbers and normal morphology. Notably, despite the inherent challenges of iSCNT, such as nuclear-cytoplasmic incompatibility, mitochondrial heterogeneity, and epigenetic reprogramming barriers^[14]^, our study successfully generated well-developed blastocysts. Immunofluorescence staining further confirmed that these interspecies cloned embryos possess normal differentiation ability.

As expected, the birth of a cloned golden wild yak was completed at Xizang Dangxiong Yak Breeding Innovation Base (4200 m.a.s.l.). Notably, previous interspecies cloning studies on yaks had only achieved early pregnancy in surrogate cows, with the fetus dying at 120 days^[15]^, indicating that maintaining interspecies cloned pregnancies to full term in yaks faces long-term challenges. Certainly, the failure of pregnancy following the transfer of yak embryos into cattle might also be attributed to the disparity in gestation length between the two species. To overcame the obstacles of embryonic arrest and miscarriage encountered in previous interspecies cloning studies, the present study using domestic yaks as surrogate recipients and successfully delivered a cloned golden wild yak, demonstrating the feasibility of iSCNT technology for the breeding and conservation of golden wild yak genetic resources. Possibly, mitochondrial heterogeneity is not an insurmountable barrier in cloning, as evidenced by the successful derivation of cloned golden wild yaks using bovine oocytes as recipients and golden wild yak fibroblasts as donors. At the less, it seems that mitochondrial compatibility between cattle and yak did not preclude successful development, although the overall efficiency (both blastocyst formation rate and birth rate of cloned individual) in the present study remains low. Although the successful cloning of the golden wild yak has expanded the applications of iSCNT technology, there are also issues that need to be explored. This study only obtained one live cloned individual, indicating that cloning efficiency still needs improvement. At the same time, reducing the miscarriage rate during pregnancy is also a future research direction. What’s more, further investigation should reveal the mechanisms of epigenetics in interspecies cloning and the effects of the surrogate mothers’ intrauterine environment on cloned embryos, to improve cloning efficiency and the survival rate of cloned animals.

## 4. Materials and methods

### 4.1 Ethics and animals

This study was approved and monitored by the Animal Welfare and Research Ethics Committee at the Institute of Animal Sciences, Chinese Academy of Agricultural Sciences (No. IAS2024-162). At the same time, all animal experiments were conducted in accordance with the guidelines established by this committee.

### 4.2 Donor cells collection and culture

The donor cells used in this study were derived from the ear tissue of a childhood male golden wild yak from Xizang Geye Wildlife Rescue Station (4800 m.a.s.l.). The biopsy was washed in phosphate-buffered saline (PBS), stored at low temperature, and transported back to the laboratory within 8 hours. The tissue was rewashed with 75 % ethanol and PBS. Subsequently, the epidermal hair and cartilage at the ear edge were removed. The tissues were enzymatically digested with 0.25 % trypsin for 30 minutes. The cell pellet was collected in Dulbecco’s Modified Eagle Medium (DMEM) supplemented with 10 % fetal bovine serum (FBS) after digestion termination, and then plated into a 10 cm culture dish under a humidified 5 % CO2, 38.5 °C incubator. After 7 days in culture, the fibroblasts reached 80 % to 90 % confluence. Then they were passaged by using 0.25 % trypsin.

### 4.3 Preparation of recipient cells

Ovaries were collected from a local slaughterhouse at Beijing that primarily processed beef and dairy cattle, and transported to the laboratory in physiological saline within 3-4 hours. Subsequently, they were disinfected with 75 % ethanol and washed 2-3 times with physiological saline. 2-6 mm follicles were aspirated using a 10 mL syringe. Then, collect and select the cumulus-oocyte complex (COCs) from these follicles. The COCs were transferred to *in vitro* oocyte maturation medium and cultured at 38.5 °C with 5 % CO_2_ and saturated humidity environment for 16-18 hours.

### 4.4 Production of iSCNT embryos

The first polar body and surrounding cytoplasm were removed from the matured oocyte. Donor cells were transferred to the perivitelline space of the enucleated oocyte. Following transfer, the cell-cytoplast couplets were induced to fuse by an ECM2001 Electrocell Manipulator in the fusion solution. The parameters of electric fusion were 1.25 kV/cm, 15 μs, two pulses of direct current. Fused embryos were treated with 5 µmol/L ionomycin for 5 minutes, and subsequently activated in 1.9 mmol/L 6-dimethylaminopurine for 4 hours. Then, iSCNT embryos were cultured further in CR1aa media at 38.5 °C in a humidified 5 % CO_2_ atmosphere for the assessment of blastocyst development.

### 4.5 Blastocyst vitrification

A 0.25 mL plastic straw (006452, imv, France) was gently heated to soften, then pulled and stretched to form an open-pulled straw (OPS). The OPS was prepared to specifications with an outer diameter of 0.22 ± 0.01 mm and a wall thickness of 0.02 mm. After cooling in air, the narrow end of the OPS was cut with a surgical blade to a length of 2.5 ± 0.5 cm.

Blastocysts were transferred from culture medium into a basal medium (HEPES-buffered M199 supplemented with 20% fetal bovine serum (FBS)) and equilibrated for 5 minutes. After being loaded into the OPS, blastocysts were transferred into an equilibration solution, in which basal medium was supplemented with 7.5% dimethyl sulfoxide (DMSO) and 7.5% ethylene glycol (EG), and equilibrated for 5 - 6 minutes. The blastocysts were then moved into a vitrification solution (basal medium supplemented with 15% DMSO, 15% EG, 0.5 M trehalose, and 1% polyvinylpyrrolidone) and held for 30 seconds. Immediately after loading embryos into the OPS, the straw was plunged into liquid nitrogen for cryopreservation.

### 4.6 Thawing and embryo transfer

The OPS was removed from liquid nitrogen, and the narrow end of the OPS was immediately immersed in a petri dish containing 1 mol/L trehalose thawing solution pre-warmed on a heating stage (38–39 °C). After the solution melted, blastocysts were released from the OPS and gently mixed by shaking, followed by equilibration for 1 minutes. The blastocysts were then sequentially transferred into 0.5 mol/L trehalose thawing solution for 3 minutes, 0.25 mol/L trehalose thawing solution for 3 minutes, and basal medium for 5 minutes. Finally, embryos were transferred into Holding medium and maintained until embryo transfer.

Adult domestic yaks were induced to synchronized oestrus and selected as surrogates. Well-developed blastocysts were transferred into the uteri of these yaks. Pregnancies were confirmed at day 60 by ultrasound examination.

### 4.7 Cell viability and cell apoptosis assay

Cell viability was determined using the CCK-8. P5 cells in good growth condition were selected and seeded at approximately 2000 cells per well in a 96-well plate. After 48 hours, the triplicate wells were chosen for the assay, and 10 μL CCK-8 solution was added to each well, followed by incubation in the incubator for 2 hours. Subsequently, the absorbance at 450 nm was measured using the microplate reader. Measurements were performed consecutively for 7 days.

Cell apoptosis analysis was performed according to the Annexin V-FITC Apoptosis Detection Kit. P5 cells were collected, washed with PBS, and then resuspended in 1 × Binding Buffer. 5 μL Annexin V-FITC and 5 μL PI were added to the suspended solution. Following the mix and place at 37 °C for 15 min, cell apoptosis was assessed by a flow cytometer.

### 4.8 Karyotype Analysis

The fibroblast cells were cultured in medium containing 0.075 μg/mL colchicine for 2 hours. Cells were digested with EDTA-free trypsin, followed by the addition of pre-warmed 0.075 mol/L KCl and incubation at 37 °C for 50 min. The fixed solution (methanol: acetic acid = 3:1) was added to the cells for 20 minutes. Repeat. Cells were resuspended in a small volume of fixed solution, dropped onto a slide, and stained with Giemsa for 10-15 minutes. Under a microscope, the number and morphology of chromosomes were identified and photographed.

### 4.9 Immunofluorescence staining in embryos

Embryos were fixed with 4 % paraformaldehyde in 1 × PBS for 35 minutes, permeabilized with 0.5 % Triton X-100 for 35 minutes, and blocked with 1 % BSA for 6 hours. Subsequently, the samples were incubated with diluted primary antibodies (CDX2; SOX2) at 4 °C overnight, followed by washing in 1 × PBS, and then incubated with diluted secondary antibodies at 37 °C for 60 minutes in the dark. Finally, the brains were mounted using Vectashield DAPI (H-1200) mounting media and observed under a laser scanning confocal microscope.

### 4.10 Short tandem repeats (STR) analysis

Ear tissues were collected from the cloned golden wild yak, donor cells, and surrogate yaks, followed by their genomic DNA. Multiple cattle-specific STR loci were amplified by multiplex fluorescent PCR. The size of the amplified products was detected by capillary electrophoresis to verify the genetic origin of the cloned individual.

### 4.11 Statistical analysis

All data were analyzed using IBM SPSS 22.0 statistical software for one-way analysis of variance. All experiments were repeated at least three times. Data are presented as mean ± S.E.M. *P* < 0.05 is considered statistically significant.

## Acknowledgement

This work was supported by the National Key R&D Program of China (2024YFD1200700), Science and Technology Project of Xizang Autonomous Region (grant number GZKJ-003, XZ202301ZY0003N, XZ202501ZY0031, XZ202402ZY0012), and Innovation Program of Chinese Academy of Agricultural Sciences (CAAS-CSAB-202402).

## Author contributions

D.Y., D.L., Y.H. and W.B. provided the conceptualization, funding acquisition, writing review and supervision. D.Y., Q.Z., L.C., S.G., Y.Z., K.Y., J.W., B.P., Y.L., and G.Z. and conducted investigation and formal analysis. C.L. and Y.H. conducted the literature research and wrote the original draft. All authors have read and approved to publish the article.

